# Namco: A microbiome explorer

**DOI:** 10.1101/2021.12.15.471754

**Authors:** Alexander Dietrich, Monica Steffi Matchado, Maximilian Zwiebel, Benjamin Ölke, Michael Lauber, Ilias Lagkouvardos, Jan Baumbach, Dirk Haller, Beate Brandl, Thomas Skurk, Hans Hauner, Sandra Reitmeier, Markus List

**Author notes:** joint last author (Sandra Reitmeier, Markus List). joint first author (Alexander Dietrich, Monica Steffi Matchado). Corresponding authors: Markus List and Sandra Reitmeier.

## Abstract

16S rRNA gene profiling is currently the most widely used technique in microbiome research and allows for studying microbial diversity, taxonomic profiling, phylogenetics, functional and network analysis. While a plethora of tools have been developed for the analysis of 16S rRNA gene data, only a few platforms offer a user-friendly interface and none comprehensively covers the whole analysis pipeline from raw data processing down to complex analysis. We introduce Namco, an R shiny application that offers a streamlined interface and serves as a one-stop solution for microbiome analysis. We demonstrate Namco’s capabilities by studying the association between a rich fibre diet and the gut microbiota composition. Namco helped to prove the hypothesis that butyrate-producing bacteria are prompted by fibre-enriched intervention. Namco provides a broad range of features from raw data processing and basic statistics down to machine learning and network analysis, thus covering complex data analysis tasks that are not comprehensively covered elsewhere. Namco is freely available at https://exbio.wzw.tum.de/Namco/.

**Impact statement:** Amplicon sequencing is a key technology of microbiome research and has yielded many insights into the complexity and diversity of microbiota. To fully leverage these data, a wide range of tools have been developed for raw data processing, normalization, statistical analysis and visualization. These tools are mostly available as R packages but cannot be easily linked in an automated pipeline due to the heterogeneous characteristics of microbiome data. Instead, user-friendly tools for explorative analysis are needed to give biomedical researchers without experience in scripting languages the possibility to fully exploit their data. Several tools for microbiome data analysis have been proposed in recent years which cover a broad range of functionality but few offer a user-friendly and beginner-friendly interface while covering the entire value whole value chain from raw data processing down to complex analysis. With Namco(https://exbio.wzw.tum.de/namco/), we present a beginner-friendly one-stop solution for microbiome analysis that covers upstream analyses like raw data processing, taxonomic binning and downstream analyses like basic statistics, machine learning and network analysis, among other features.

## Introduction

Over the past decade, microbiome research has contributed to our understanding of human health and microbiome-associated diseases with implications for diagnosis, prevention and treatment (1). Several studies have discovered important links between the gut microbiome and human diseases including diabetes (2), cancer (3), inflammatory bowel disease (4) and brain disorders (5).

Currently, microbiome datasets are generated using either targeted (amplicon) gene sequencing to characterize the microbial composition and phylogeny or shotgun metagenomics to study, in addition, the gene-coding and functional potential of the microbiome. Due to the comparably lower sequencing costs, sequencing of the 16S rRNA gene is the most commonly available method. Analysis of 16S (or 18S in the case of eukaryotes) rRNA genes can be grouped into four steps. First, amplicons are clustered into operational taxonomic units (OTUs) (6) or amplicon sequence variants (ASVs) (7). For OTUs, sequencing errors are addressed by choosing a similarity threshold of typically 97% for clustering whereas the latter employs a denoising strategy to identify the unique error-corrected sequence of an organism (8). Several benchmark studies proved that denoising methods provide better resolution and accuracy than OTU clustering methods (7, 9). Along with taxonomic profiling, various measures of alpha and beta diversity are typically computed to study the microbial diversity within and between samples, respectively. Additional analysis steps include, for instance, (i) differential abundance analysis to identify bacteria and/or functions that differ between groups of interest, (ii) in silico inference of metagenomes for functional profiling and (iii) microbial association analysis through correlation, co-occurrence and network inference methods.

Many open-source tools have been developed for microbiome data analysis (Table 1). *Mothur (10), QIIME2 (11), DADA2 (8)* and *LotuS2* (12) offer processing of raw sequencing files through clustering and annotation of 16S rRNA genes and provide OTU or ASV tables which serve as an input for further downstream analysis. The *QIIME2 (11)* pipeline offers more than 20 plugins for downstream analysis including q2-sample-classifier for supervised classification and regression analysis (13), q2-longitudinal for time-series analysis (14), and plugins for compositional data analysis (15). In addition, a plethora of R packages have been implemented to perform statistical analysis and high quality visualizations on amplicon tables, like *Phyloseq (16, 17)* and *themetagenomics (18)*. Even though R scripts leveraging these tools offer a powerful approach to analyze microbial data, their application can be demanding for users without scripting knowledge and bioinformatics training. Hence, there is a need for user-friendly tools that can support end-to-end exploration of microbiomes. To this end, several web-based tools have been developed including *microbiomanalyst (19), IMNGS (20), iMAP (21), MGRAST (22), wiSDOM (23), VAMPS (24), Shiny-phyloseq (17)* etc. However, most of these tools (i) cover only the downstream part of the analysis and often omit raw data processing, (ii) offer only standard analysis and are thus not sufficient for more complex data sets, (iii) omit functional profiling or use outdated approaches such as *Tax4FUN (25)* and *PICRSUt1 (26)* which were outperformed by *PICRUSt2 (27)*, (iv) do not offer confounder analysis (v) offer no support for time-series and (vi) lack support for machine learning and (vii) do not construct microbial association networks and differential networks on different taxonomic ranks.

**Table 1:**
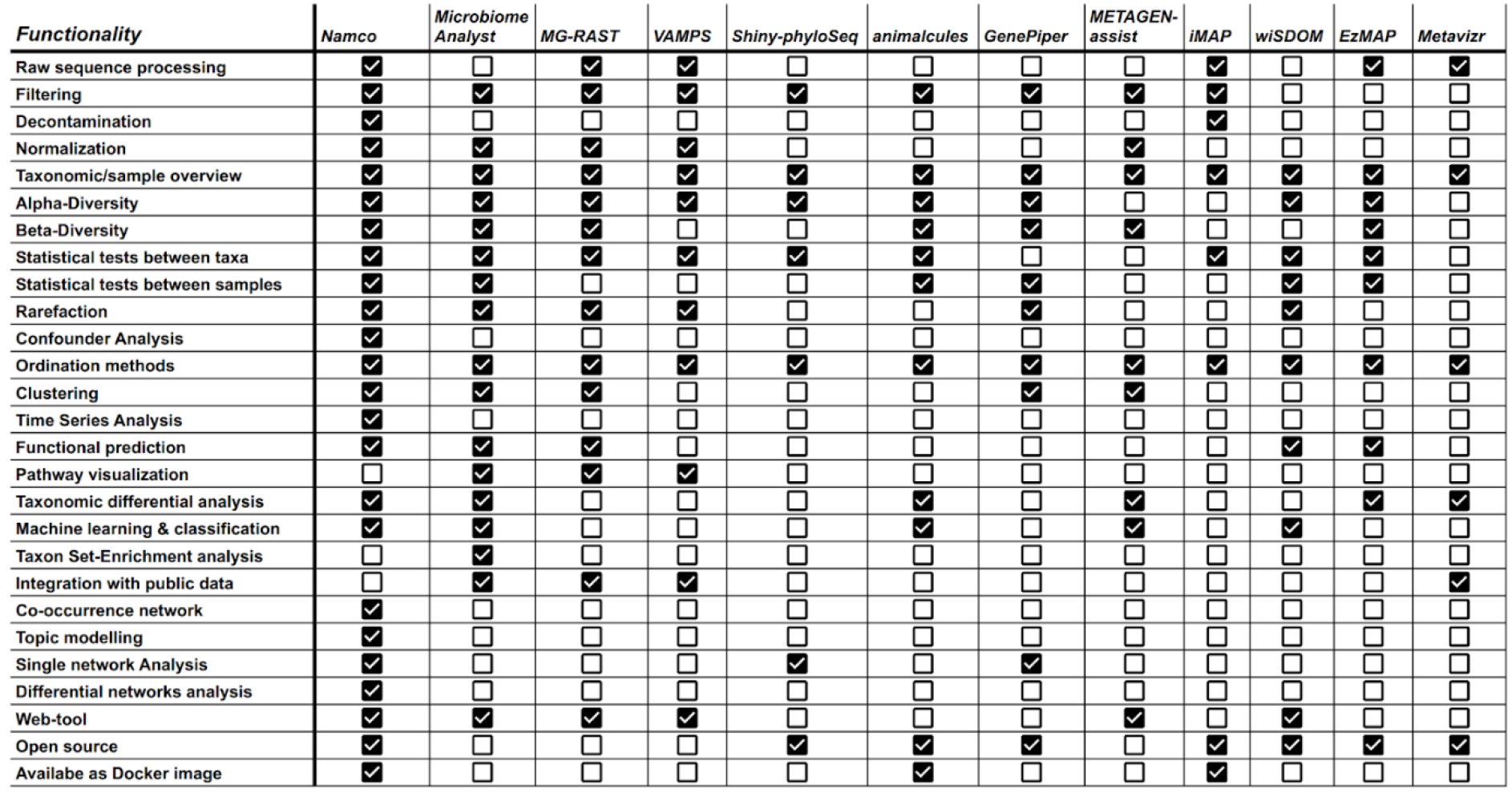
Comparisons of Namco with other web-based tools for microbiome data analysis.

To address these limitations, we introduce Namco, an R (v 4.1) shiny application that offers a streamlined and user-friendly interface and serves as a one-stop solution for microbiome analysis. Namco provides a broad range of features from raw data processing and basic statistics down to machine learning and network analysis, thus covering complex data analysis tasks that are not comprehensively covered elsewhere (see Table 1 for a comparison with other tools). Namco’s thoroughly documented and easy-to-use graphical interface is intended to eliminate the use of command-line arguments during the data processing, making advanced microbiome analysis accessible to a broad range of biomedical researchers. Privacy legislation such as the European Union’s General Data Protection Regulation (GDPR) can prevent users from uploading their data to web tools such as Namco. Hence, we make Namco available under an open source license and release it as a Docker container, which can be executed locally (e.g. using Docker Desktop) or can be safely deployed on a local protected server.

## Methods

### Input file formats

For raw data processing, Namco accepts both single and paired-end FASTQ files which are processed internally based on DADA2 (8) or LotuS2 (12). Alternatively, users can start their analysis with a previously generated OTU or ASV table which can either include relative abundance values or read counts. For simplicity, we refer to these as features throughout the manuscript. Features should be labelled either by their taxonomy or alternatively by a unique id which Namco can map to a separate tabular file with taxonomic labels that can be uploaded separately by the user. Optionally, users can upload a metadata file containing additional information for each sample which can be leveraged for groupwise differences, correlation, confounder or condition-specific analysis, for instance. Finally, users can optionally provide a tree file in newick format for phylogenetic tree analysis, diversity analysis or to create ecologically-organized heatmaps. The entire workflow of Namco is presented in Figure 1.

**Figure 1:**
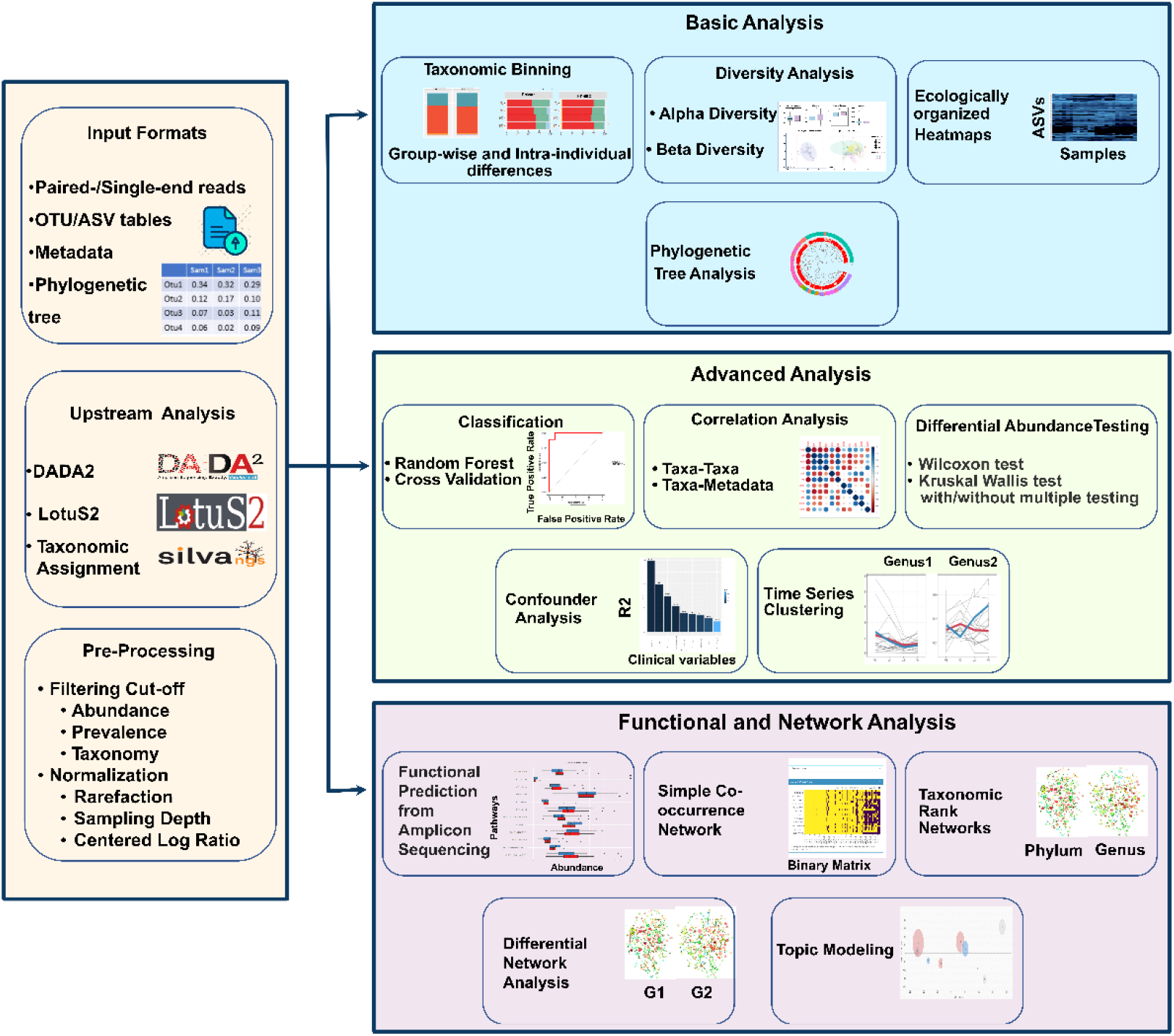
Overall workflow of Namco. Namco provides a comprehensive end-to-end analysis of microbiome data including raw FASTQ processing and filtering, down to statistical, functional and network analysis. It also provides various tables and visualization options and allows users to navigate through different data analysis tasks.

### Denoising and taxonomic assignment

Namco accepts both Ilumina single- and paired-end reads in FASTQ format. It supports two pipelines for upstream analysis, DADA2 (8) and LotuS2 (12). The latter is a user-friendly pipeline providing access to six different sequence clustering algorithms including USEARCH (6), UNOISE3 (28) and DADA2 (8). It features 21 quality filtering metrics (12) which improved the quality and consistency of the output. Users should be aware of the filtering techniques used in LotuS2 before applying these to their own data. In addition, Namco also provides a standard alone DADA2 option for denoising steps. Users can change the default parameters such as trimming length depending on the amplicon length and the quality score threshold. While filtering primer regions, the DADA2 option in Namco expects that the primer sequences are present at the start of the input reads with a constant length. The SILVA (v. 138) (29) database is used as a reference for taxonomic classification. Namco stores the *LotuS2/DADA2* output as a phyloseq object using the *phyloseq* R (16) package and passes it down to further downstream analysis. Alternatively, users can perform their own upstream process elsewhere and upload abundance tables and metadata as inputs to Namco.

### Data overview and filtering

The data overview section in Namco summarizes sample details, total number of features identified during the denoising step, and the number of groups provided by the metadata file. This gives an overall picture of the input data and will help in preparing the input data for further processing. Normalization and filtering are considered crucial steps in microbial analyses (30, 31). By default, Namco applies 0.25% of the abundance filter on the DADA2-generated features and normalizes abundance to 10,000 reads before downstream analysis. Filtering the abundance of both OTUs and ASVs > 0.25% (32) was identified as an effective threshold to prevent the identification of spurious taxa to a large extent. Alternatively, users can also choose different normalization methods and filtering percentages such as sampling depth, rarefaction and centered log-ratio transformation (33). In addition to the normalization methods, Namco offers different filtering options based on sample prevalence and relative abundance as microbiome data are very sparse in nature and often have zero counts in most samples. These rare taxa are caused by sequencing artefacts, contamination and/or sequencing errors (32). In addition, Namco utilises the decontam R package (34) which can differentiate contaminants from non-contaminants across diverse studies (35, 36) and hence improves the quality of biological conclusions in microbiome studies.

### Basic analysis

#### Visualization of taxonomic binning and ecologically-organized heatmap

Namco integrates R-scripts from *Rhea (37)* and *phyloseq (16)* to perform taxonomic profiling and diversity analysis and provides different options to visualize the distribution of dominant taxa at different ranks (domain, phylum, class, order, family and genus) among groups using barplots. In addition, taxonomic distribution can also be inferred based on intra-individual differences by visualizing taxa for individual samples. Users can download feature tables of relative abundances aggregated at different taxonomic levels and export any of the generated plots. The advanced heatmap option in Namco creates a heatmap using ordination methods such to organize the rows and columns instead of hierarchical clustering approach which gives an overview of the abundance of features across sample groups that are very high /low abundant.

#### Diversity analysis

Alpha diversity quantifies the diversity of the microbiome within a group. Namco supports five common alpha diversity measures, namely *Shannon entropy (38) and Simpson index (39)* together with their counterparts accounting for the effective number of species (40) as well as richness. Users can select different categories from the metadata to visualize alpha diversity and determine significant differences via a Wilcoxon test. Beta diversity analysis explains the variation between groups and relies on a phylogenetic tree as input along with the feature table to calculate dissimilarity. Namco supports the most common distance metrics including *weighted* and *unweighted unifrac distances, generalized unifrac (41), variance adjusted unifrac distance* and *Bray Curtis dissimilarity (42)*. Calculation of *unifrac distances* is only possible if a tree file was uploaded. The results are presented using non-metric multidimensional scaling (NMDS) and PCoA (43). In addition, it is also possible to visualize the distance as a hierarchically clustered dendrogram which helps to identify closely related samples. Significance between groups is determined by a permutational multivariate analysis of variances using the adonis function of the vegan R-package (44). P-values are corrected for multiple testing following Benjamini-Hochberg (BH) (45).

### Differential analysis

#### Differential abundance testing using simple statistical tests and association analysis

A key aim of microbial research is to identify differences in microbial composition between conditions or phenotypes. Namco reports statistically significant features between the sample groups using the non-parametric Wilcoxon test (*SIAMCAT* R-package (46) which was shown to reliably control the false discovery rate in differential abundance analysis (47). Users can choose groups that should be compared against each other and adjust the significance level as well as other filtering parameters. Differential abundance can be calculated at different taxonomic levels such as phylum or genus, where Namco aggregates the feature table accordingly. Namco shows the distribution of microbial relative abundance along with the significance and a generalized fold change (48) as a nonparametric measure of effect size. In addition to the Wilcoxon test, Namco also offers Kruskal-Wallis test to find significant differences between more than two groups.

#### Correlation analysis

Namco provides correlation analysis to reveal significant associations between taxa or between taxa and metadata such as continuous experimental variables. Namco further considers relative abundances of features at different levels (phylum, class, order etc).

#### Topic modeling

Topic modeling was originally designed to uncover hidden thematic structures in document collections (49). This concept was adapted to metagenomic analyses to explore co-occurring taxa as topics and to find topics associated with provided sample metadata (50). Namco employs the *themetagenomic R (50)* package to predict topics and to study their association with sample metadata which can be continuous, binary, categorical, or factor covariates.

#### Functional profiling

Microbial composition varies widely between individuals, making the robust identification of phenotype-associated microbial features challenging. One can hypothesize that the functional potential of the microbiome is more robustly associated with a phenotype than the microbial composition. While investigating the functional potential of the microbiome is not directly feasible with 16S rRNA gene sequencing data, several tools have been proposed for inferring the functional profile with the help of reference sequencing databases. To this end, Namco adapts the *PICRUSt2 (27)* approach, which showed improved accuracy and flexibility compared to related tools including *PICRUSt, Tax4FUN2, Piphillin* and *PanFP (27)*. Namco also performs differential analysis on the predicted KEGG orthologs, enzyme classification numbers and pathways using *Aldex2 (51)*, which was recently reported to perform best for this type of analysis **(52).** The relative abundances of significant KEGG annotation terms are plotted in a barplot along with the p-value.

#### Phylogenetic tree analysis

Phylogenetic analysis belongs to the basic steps to get an overview of the evolutionary relations between features. Namco displays the provided/calculated phylogenetic tree in circular or rectangular format. In addition, users can add two heatmap layers as taxonomic ranks and/or a meta-group. The meta-group heatmaps are colored by the abundance of the features in the corresponding meta-group.

#### Network analysis

Several methods for inferring and analysing microbial co-occurrence networks were developed to study the role of microbial interactions in association with the host (53, 54). Namco implements multiple strategies for network construction. As a simple approach, the feature abundance matrix is converted into a binary indicator matrix using an abundance cutoff (presence/absence) (default cut-off is 1: all features with a value less than 1 are considered absent, while the rest is considered present.) This cutoff can be adapted manually to get a more strict binary representation of the abundance matrix. Next, the number of co-occurring feature pair is counted across samples for each groups (e.g. case and control-pairwise) and the difference in co-occurrence counts, as well as the log2 fold-change between the groups, are calculated and displayed as a network where nodes represent features and edges represent frequent group-specific interactions. For more advanced approaches, Namco employs the *NetComi* (55) R package, where users can build microbial association networks at different taxonomy levels using nine different network construction algorithms. In addition, *NetComi* offers a method for differential network analysis between two conditions to identify pairs of taxa differentially associated between two groups.

#### Confounder analysis and explained variation

Confounding variables may mask the actual relationship between the dependent and independent variables in a study (56). Especially microbiome composition is associated with several host variables including body mass index (BMI), sex, age and geographical location, among others (57). Namco utilizes the permutational multivariate analysis of variances (adonis function of the *vegan R*-package) (44) to rule out confounding factors using available information from the user-provided metadata table. The explained variation of covariates is determined by R^2^ values which are considered significant at a p-value ≤ 0.05.

#### Classification based on random forest

Beyond differential abundance analysis, an important question is if a classification model can be trained on a minimal set of features to robustly predict the outcome (e.g. disease state or treatment response). Such models highlight the potential of microbiome data for prognostic and diagnostic purposes through biomarkers and surrogate endpoints. Namco allows users to build classification models and identify important features using machine learning algorithms. Within Namco, random forest (*ranger (58)* R package) is used as a classification tool, since it has shown good performance even on comparably small sample sizes in microbial data analysis (59, 60). By default, Namco splits the data into training and test sets and performs 10-fold repeated cross-validation. Experienced users can modify advanced parameters such as the ratio of training and test sets, the number of cross-validation folds, the resampling method and the number of decision trees. The results are summarized in a confusion matrix and a receiver-operator-characteristic (ROC) plot which helps in evaluating the model performance. The most informative features that were used for classification can be extracted as biomarker candidates for hypothesis generation and further research.

#### Time series analysis and clustering

Time-series analysis in Namco allows users to determine how microbial communities including taxa, OTU / ASV, and other features like richness change over time. For instance time-series analysis helps to study the microbial changes in response to a treatment over multiple timepoints or during different stages of host development. Namco offers different options to modify the inputs for time series line charts including displays of changes in either relative abundance or absolute abundance or richness.

#### Use Case

To illustrate the broad utility of Namco, we analyzed human fecal samples from an interventional cross-over study. The study’s aim was to develop healthier convenience food products with a increased fibre content and to foster customer acceptance of such products. Here, we analyzed if the stool microbiota is altered by the fibre-rich diet.

#### Ethics Statement

The study protocol was approved by the ethical committee of the Faculty of Medicine of the Technical University of Munich in Germany (approval no. 529/16S). The guidelines of the International Conference on Harmonization of Good Clinical Practice and the World Medical Association Declaration of Helsinki (in the revised version of Fortaleza, Brazil 2013) were considered. All study participants have given written informed consent. The study was registered at the German Clinical Trial Register (DRKS00011526).

#### Study design

The human intervention study was a single-blinded, controlled cross-over study. Volunteers were recruited from a cohort of middle-aged subjects who were broadly phenotyped within the *enable* nutrition cluster, 50% were male and 50% female. Inclusion criteria: 40-65 years old volunteers with an elevated waist circumference. For detailed information on inclusion and exclusion criteria of the enable cohort see Brandl et al (61). Study participants were invited four times to the study center. During the first visit, baseline characteristics were collected. In general, the intervention of meatloaf in a bun and pizza was performed as described in Rennekamp et al (62).

#### Phenotypic characteristics of the study group

The study group was age and sex-matched (N = 11 females, N = 10 males) and received the same intervention and placebo (Table 2). Baseline measurements were performed in the morning after an overnight fasting. Body composition and body weight were measured by using a Seca Medical Body Composition Analyser, mBCA 515, Seca GmbH & Co. KG, Hamburg, Germany. Body height was measured in a standing position without shoes using a stadiometer (Seca GmbH & Co. KG, Hamburg, Germany). BMI was calculated as weight (kg)/height (m2). Waist circumference was measured at the midpoint between the lowest rib and the iliac crest with a measuring tape (Seca GmbH & Co. KG, Hamburg, Germany).

**Table 2.** Overview of the study group characteristics. Indication of the mean values and the standard deviation for the participants is given, besides significant difference in traits between the sexes.

|  | Mean | SD | Differences between sexes |
| --- | --- | --- | --- |
| Weight [kg] | 90.14 | 11.42 | 0.0080 (**) |
| Height [m] | 1.73 | 0.08 | 1.35e-05 (***) |
| BMI [kg/m<sup>2</sup>] | 30.12 | 2.41 | 0.8490 (ns) |
| Fat-free mass [%] | 62.98 | 6.74 | 2.40e-08 (***) |
| Fat mass [%] | 37.02 | 6.74 | 2.40e-08 (***) |
| Skeletal muscle mass [kg] | 27.55 | 6.28 | 9.34e-07 (***) |
| Visceral fat [kg] | 3.24 | 1.32 | 4.55e-05 (***) |
| Waist circumference [cm] | 101.3 | 7.26 | 0.0058 (**) |

#### Sample preparation

The participants were asked to visit the study center sober (10 hours before the visit) and received the intervention or placebo meal in the study center. Additionally, a capsule with food coloring was administered. The intake of the dye stains the stool green which helps to associate the collected samples and food intake. The time of the meal, as well as the time of the excretion, were recorded and showed a mean transit time of 34.74 ± 24.69 h. Since a coloring capsular was administered together with the meals, the stool samples can be assigned to the meal. The dye causes a visible green coloration of the sample, and a recognizable coloration was noted in the data. Participants consumed two different types of food (meatloaf in a bun and a pizza) both, either enriched with fibre (intervention) (IM) or not (placebo) (M). The first interventional meal (meatloaf in a bun, IM1) the white bread roll in the fibre-enriched meal contained an additional 5.7% wheat fibre (VITACEL^®^ WF600) and the meatloaf (Leberkas) a mixture of 3.1% wheat fibre and 4.5% resistant dextrin. The second intervention (pizza, IM2) was also fibre-enriched containing up to 20 g fibre with 3.0 % wheat fibre, 2.4% powdered cellulose and 2.1% inulin (Table 3). The intervention meals thus constituted a major part of the recommended daily fibre intake. As the fibre content is above 6 g per 100 g, the food products are considered to count as high fibre products.

**Table 3.** Nutritional values per serving for the intervention (enriched) and the placebo (standard) meatloaf and salami pizza meal as well as the difference between intervention and placebo meal.

|  | **Portion meatloaf with bun 240 g** |  | **Portion salami pizza 320 g** |  |
| --- | --- | --- | --- | --- |
|  | **Enriched** | **Standard** | **Enriched** | **Standard** |
| Energy [kcal] | 413 | 587 | 829 | 876 |
| Fat [g] | 13 | 35 | 41 | 45 |
| Carbohydrate [g] | 47 | 47 | 75 | 83 |
| Total fibre [g] | 19 | 2.9 | 20 | 6 |

#### Sample preparation and sequencing

For the analysis of gut microbiota, the 16S rRNA gene was sequenced at the ZIEL Core Facility Microbiome, Technical University Munich, Germany. A detailed description of the samples preparation and sequencing are described elsewhere (63). Briefly, sample DNA was isolated following an in-house developed protocol. Targeting the V3V4 region of the 16S rRNA gene, samples were amplified and purified. Pooled amplicons were paired-end sequenced on an Illumina MiSeq. Sequencing data is available under BioProject ID PRJNA774891

#### Research question

An increased fibre intake was shown to be protective against the development of cardiovascular (64, 65) and malignant diseases (66, 67) and there are specific health claims associated with specific types of fibres. In this study, we examined the presence of butyrate-producing bacteria which could be promoted by the fibre-enriched intervention and thus prove that dietary aspects can have a permanent effect on the gut microbial composition.

## Results

### Diversity analysis

We studied changes in the microbial composition following dietary intervention using Namco. Paired-end FASTQ files were processed using the DADA2 denoising step embedded in Namco with the default parameters. During the DADA2 step, a 0.25% abundance-based filter was applied to reduce sparsity (32). ASVs were normalized to 10,000 reads before downstream analysis and outliers were removed. Additionally, a prevalence filter cutoff of 10% was introduced. No significant differences were observed in alpha diversity measures between the IM and M groups: Shannon, richness, Simpson Index, effective Shannon entropy or effective Simpson entropy. Likewise, no significant clustering was found in beta diversity including unweighted and weighted Unifrac between IM and M (Figure 2).

**Figure 2.**
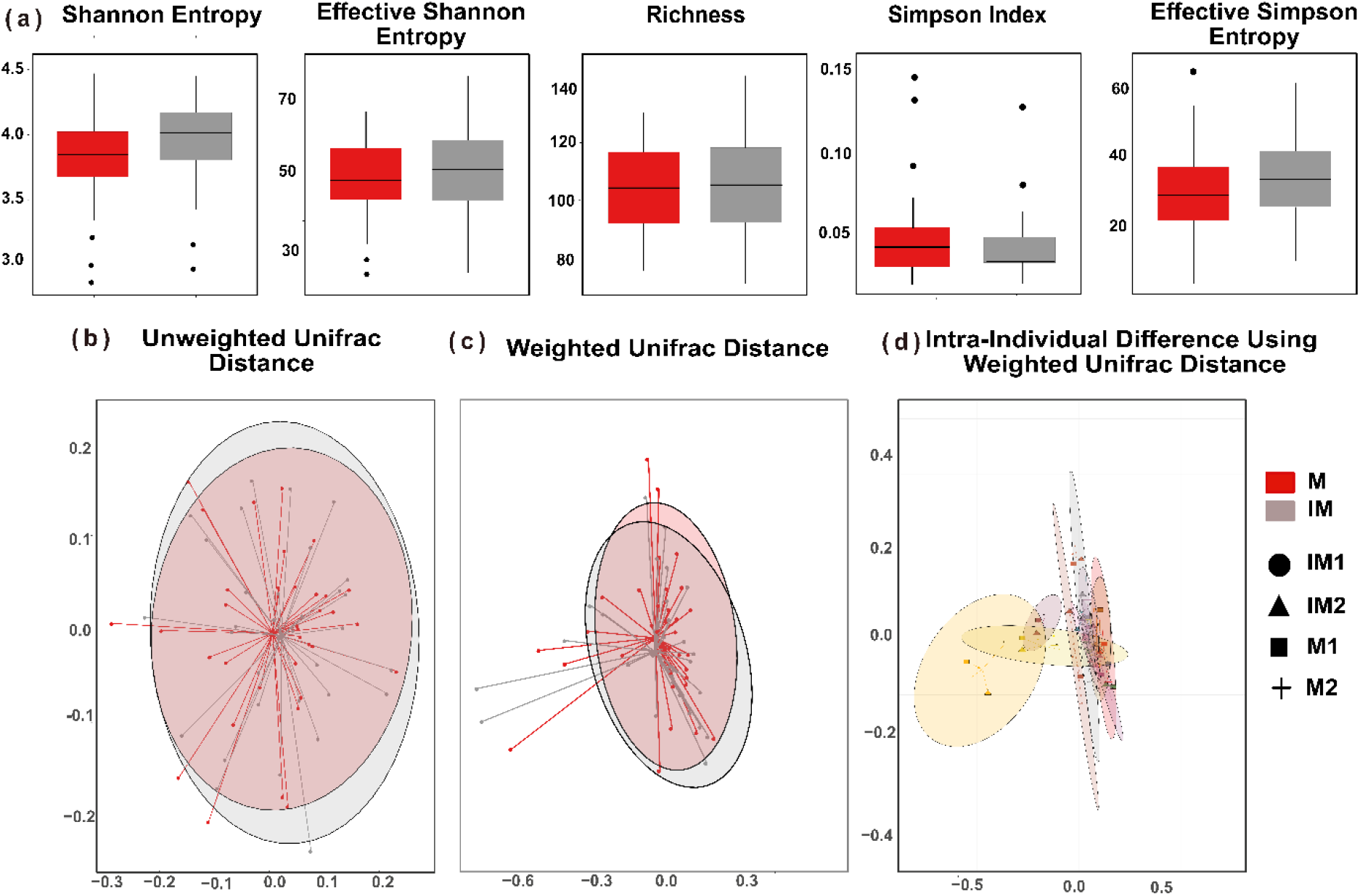
**(a):** Alpha diversity measures associated with intervention (IM) and non-intervention meals (M). There was no significant difference identified between the groups. (b) nMDS visualizations of beta diversity analysis using the unweighted or weighted (c) unifrac distance. (d) nMDS visualizations of beta diversity for intra-individual patients across two intervention meals and their respective control using weighted unifrac distance.

### Taxonomic distribution

Overall, 6 phyla and 77 genera were observed in all groups. Dominating phyla in both intervention (IM1 and IM2) and non-intervention groups (M1 and M2) were Firmicutes and Bacteroidota, contributing up to 90% to the total bacterial composition. The relative abundance of Firmicutes was found to be slightly higher in the IM group compared to the M group. Bacteroidota were slightly more abundant in the M group. Other phyla such as Actinobacteria, Verrucomicrobia and Proteobacteria showed < 5% mean relative abundance between IM and M groups **(Figure 3 (a))**. *Ruminococcaceae Incertae Sedis* was significantly different between the IM and M groups. Overall, except *Ruminococcaceae Incertae Sedis*, no other significant difference was observed at the phylum level between IM and M groups. However, intra-individual heterogeneity was observed at the abundance on phylum level **(Figure 3 (b))** at each intervention. On the phylum level, we observed differences in the relative abundance of Firmicutes and Actinobacteria from 34% to 77% and from 0.46% to 10.56%, in the IM and M, respectively. Similarly, the relative abundance of Proteobacteria also varied from 0.03% to 5.6 % and 0.07% to 8.6% in the IM and M, respectively. On the genus level, most of the individuals showed a uniform distribution except for one individual who showed a high level of *Prevotella* (54% of relative abundance).

**Figure 3:**
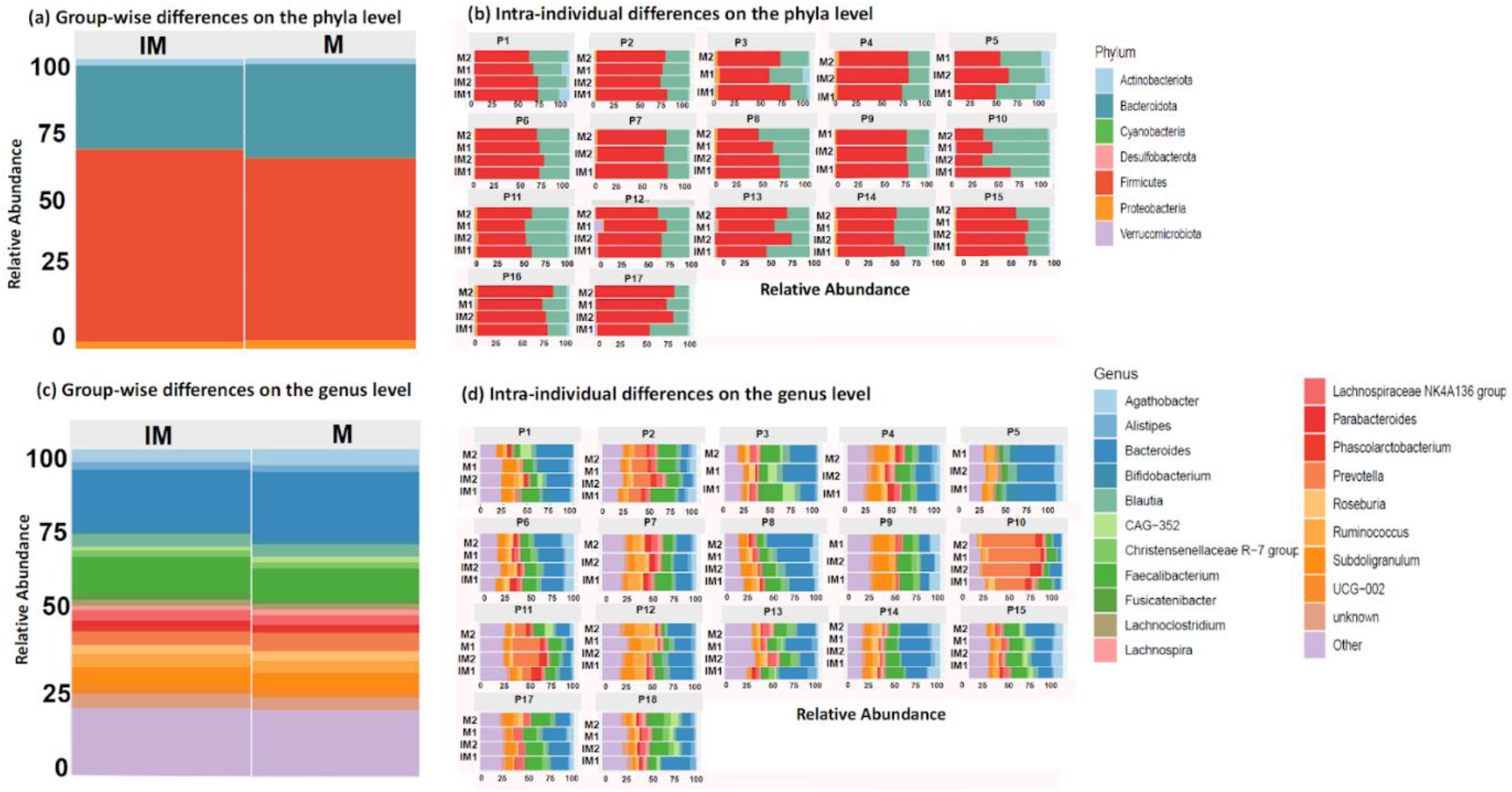
(a, c) Relative abundances of phyla and genus between intervention and non-intervention groups. (b,d) Bar plots show inter-individual variation in the gut microbiome between intervention and non-intervention meals.

The top 20 genera are shown in **Figure 3 (c)** and intra-individual differences are plotted in **Figure 3 (d)**. *Bacteroides was* the most abundant genus in both IM and M groups followed by *Faecalibacterium, Prevotella and Agathobacter*. Due to the similarity observed at the genus level between IM and M meals, a non-parametric paired Wilcoxon test was applied to the relative abundance to identify microbial changes between (IM1, IM2) and their respective controls (M1 and M2). Samples with missing information regarding what kind of meals were adminintrated were removed during the following analysis. In total, five genera: *Anaerostipes, Ruminococcaceae Incertae Sedis, Parabacteroides, Fusicatenibacter and Butyricicoccus* were significantly different in abundance prior to multiple corrections between intervention meals and normal meals. After multiple testing corrections with BH, only *Ruminococcaceae Incertae Sedis* remained significant. The relative abundance of *Anaerostipes, a* butyrate-producing bacterium, was found to be higher in IM2 compared to M2. *Anaerostipes* is a gram-positive and anaerobic bacterial from the family of *Lachnospiraceae* and found to be highly expressed in a normal healthy gut (68). As a validation, previous studies stated that the abundance of *Anaerostipes* increases with fibre-rich diets and is negatively correlated with BMI (69, 70). *Ruminococcaceae Incertae Sedis* and *Parabacteroides* also showed a significant difference between IM1 and M1. **(Figure 4)**. *Ruminococcaceae* is also known for producing short-chain fatty acids (SCFA) including butyrate which promotes a healthy bowel (71) and is nominally protective of weight gain (72). In addition, *Ruminococcus bromii* is reported as the key species in fermenting resistant starch, which in turn helps in conferring health benefits including weight control and protection against diabetes *(73). Parabacteroides* have been reported to have metabolic benefits and a negative correlation with BM I (74, 75). One of the species of *Parabacteroides (P. distasonis)* is also reported to be part of the core gut microbiome (76–78) and has the ability to produce succinic acid and of bile acid in promote the increase terms of regulating host metabolism (75, 79). Ezeji et al. also found *Parabacteroides* to be enriched in fibre-rich dietary intervention groups (80). At the genus level, dietary fibre intervention significantly promoted the growth of beneficial genera *Anaerostipes, Ruminococcaceae Incertae Sedis* and *Parabacteroides*. Additionally, *Fusicatenibacter* was found significantly higher in IM1 than M2 and *Butyricicoccus* was found higher in the M1 group compared to IM2.

**Figure 4:**
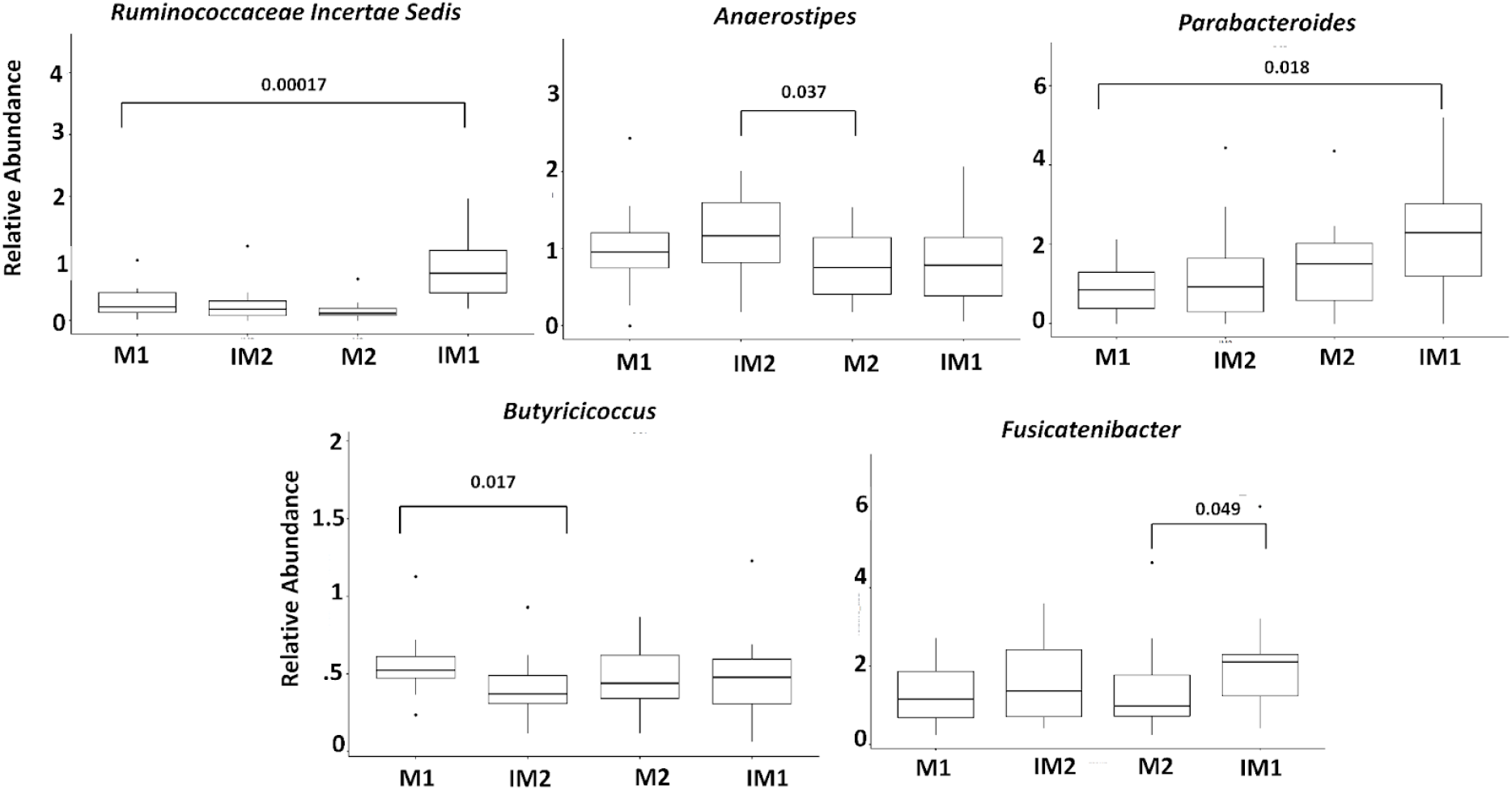
Boxplots representing the significant differences between mean proportions of genera between four meal groups. Significance was tested using non-parametric Wilcoxon Rank test. IM1 and IM2 represent the first and second interventional meals, respectively. M1 and M2 represent first and second non-intervention meals, respectively.

### Correlation of gut microbial composition and clinical metadata

To study possible associations between features and continuous variables of interest such as fat free mass [%], fat mass [%], skeletal muscle mass [kg], BMI and age, Spearman correlation was calculated at the phyla level **(Figure 5 (a))**, showing that formerly known as Deltaproteobacteria) was negatively correlated with BMI followed by Verrucomicrobiota, and Firmicutes Actinobacteriota. On the genus level **(Figure 5 (b))**, the genera *Phascolarctobacterium, Lachnospira, Lachnospiraceae FCS020 group, Prevotella, Alistipes and Oscillospiraceae UCG-005* were significantly but positively correlated with BMI, whereas *Bacteroides* and *Ruminococcus* and *Lachnoclostridium* were negatively correlated with BMI. *Prevotella, Lachnospiraceae FCS020 group* and *Phascolarctobacterium*, which were positively correlated with BMI, were also slightly less abundant in the IM group compared to the M groups. *Anaerostipes, [Eubacterium] ruminantium group* and *Lachnospira* were also positively associated with fat mass [%]. Conversely, *the Rikenellaceae* RC9 *gut group and Clostridia UCG-014* were negatively associated with fat mass percentage.

**Figure 5:**
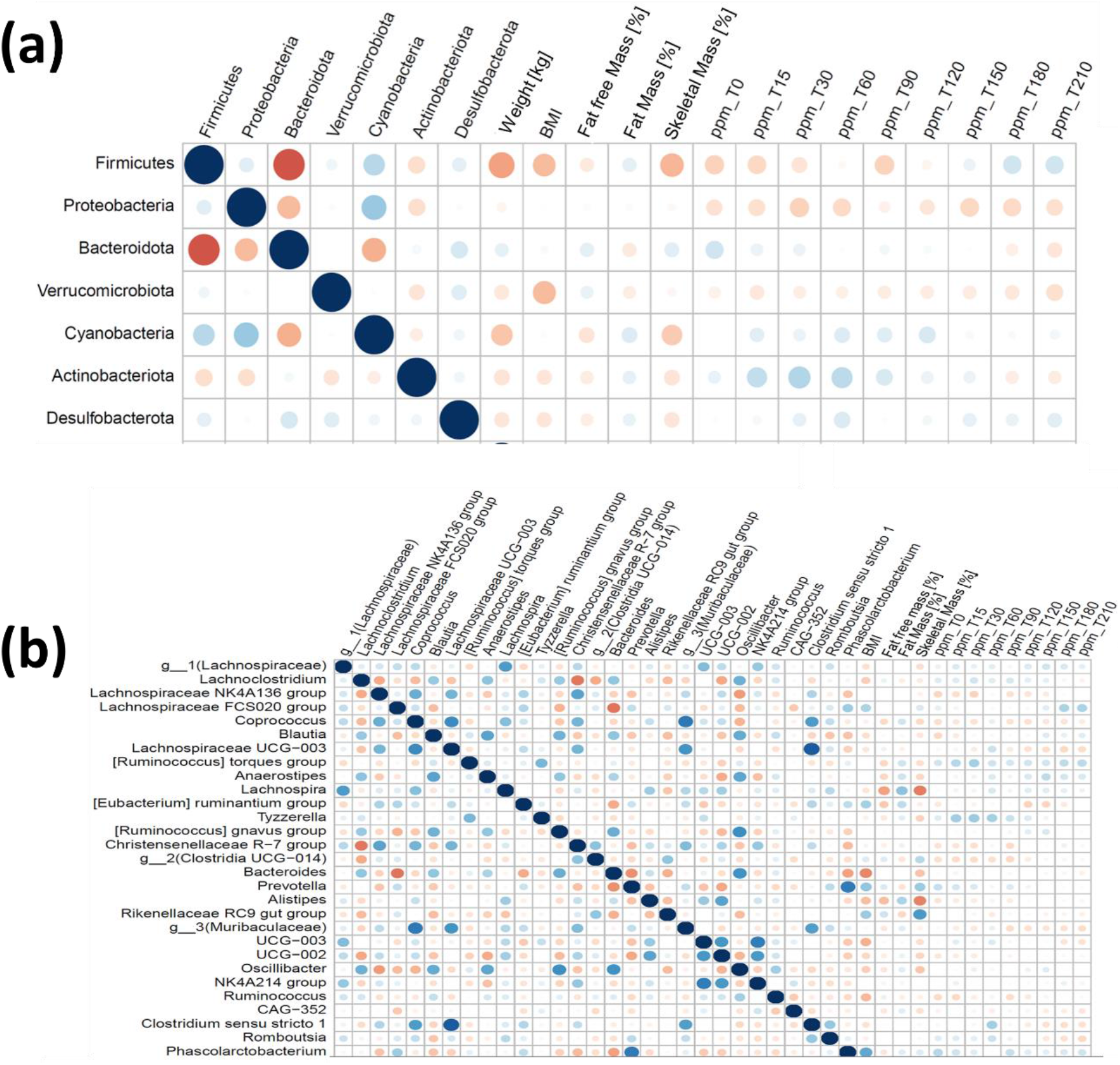
Correlation between gut microbial composition and clinical variables at phyla (a) and (b) genus level

**Figure 6:**
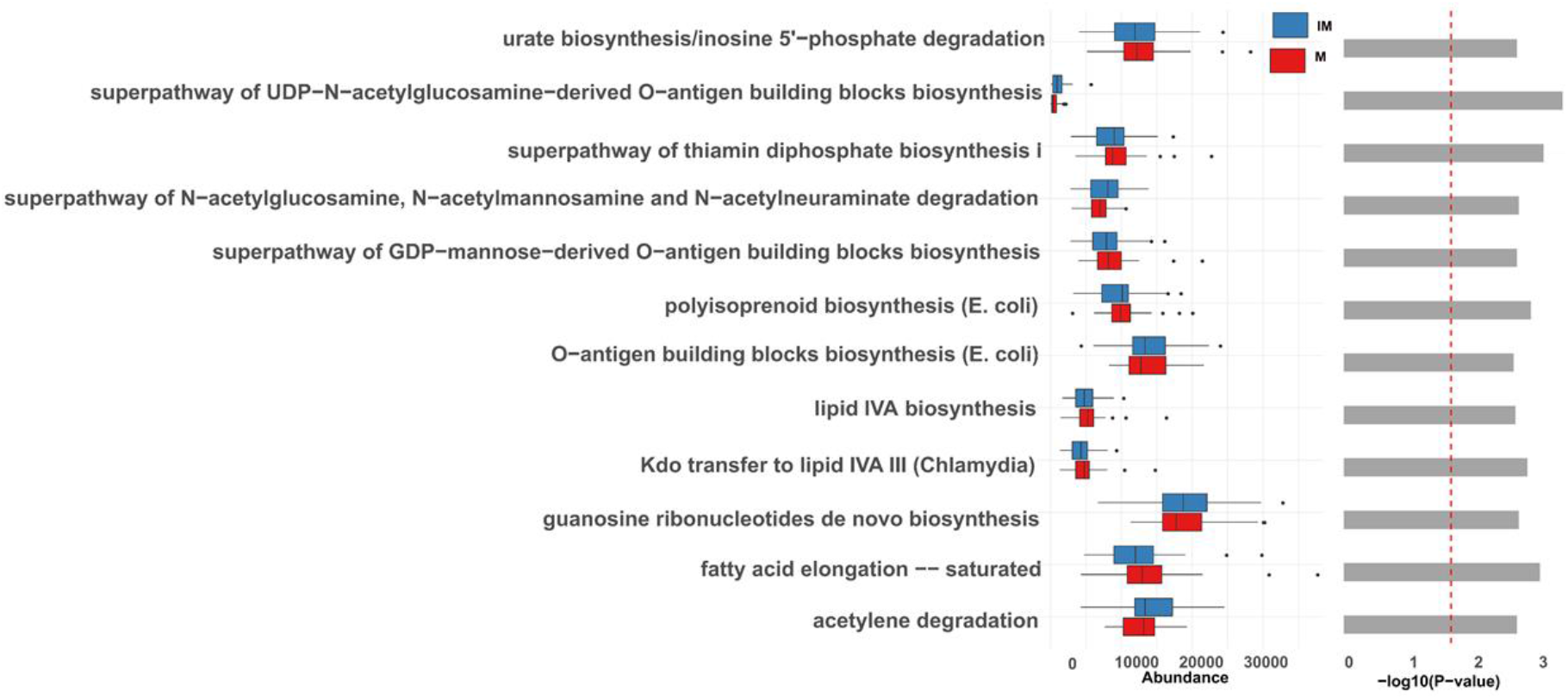
Bar plot showing pairwise-group comparison of significant pathways. Wilcoxon test was performed on relative abundance and extended error bar plots were used for the comparison between IM and M groups. Only predicted functions and pathways with p < 0.05 are shown. Bar plots on the left side display the mean proportion of each KO term with - log10 p-value while the right display the mean proportion of each KEGG pathway with - log10 p-values.

### Functional analysis

The built-in PICRUSt2 option of Namco was used to infer the functional differences of the microbial communities between the IM and M groups. Similar to differential abundance analysis at the taxa level, significant differences were calculated with a paired Wilcoxon rank test between the IM and M groups. In total, 82 KEGG Orthologues (KO) were significantly different between the groups without correction for multiple testing. Significant KO terms with p-value < 0.05 were grouped according to KO categories in order to understand their functions. The majority of the 76 KO terms belonged to metabolic categories (level 1) and were further divided into 11 sub-categories (level 2) including carbohydrate metabolism, amino acid metabolism, energy metabolism (Oxidative phosphorylation), lipid metabolism, metabolism of cofactors and vitamins and glycan biosynthesis and metabolism. Among these, the KO term K00845 (glucokinase) is part of the amino sugar and nucleotide sugar metabolism which was enriched in the IM groups but not in the M groups. Previous studies suggested that high fibre intake shows a positive impact on glucose and fat metabolism in humans (81). In support of that, ATP-binding cassette (ABC) transporters such as K02018, K10823, K15580 and K15583 were also upregulated in IM. Previous studies suggested that *Firmicutes*, which were slightly more abundant in the IM group, encode ABC transporters which belong to transport ATPase groups on the bacterial plasma membrane. These transporters, which are essential to transfer glucose to the other side of the plasma membrane (82) also help in transporting anti-inflammatory butyrate from bacterial digestion of dietary fibres (83). Differences in the abundance of ABC transporters and glucokinase are shown in Figure 7. Aspartate aminotransferase (AST) (K00812) was downregulated in the IM group compared to the M group. AST is an important biomarker for liver damage. Few studies showed that fibre-riched diets reduce the level of AST (84). We also identified twelve pathways as significantly different between IM and M groups, namely fatty acid elongation saturated (FASYN-ELONG-PWY), super pathway of N-acetylglucosamine, N-acetylmannosamine and N-acetylneuraminate degradation (GLCMANNANAUT-PWY), lipid IVA biosynthesis (NAGLIPASYN-PWY), O-antigen building blocks biosynthesis (E. coli) (OANTIGEN-PWY), acetylene degradation (P161-PWY), polyisoprenoid biosynthesis (E. coli) (POLYISOPRENSYN-PWY), urate biosynthesis/inosine 5’-phosphate degradation (PWY-5695), Kdo transfer to lipid IVA III (Chlamydia) (PWY-6467), guanosine ribonucleotides de novo biosynthesis (PWY-7221), super pathway of GDP-mannose-derived O-antigen building blocks biosynthesis (PWY-7323), super pathway of UDP-N-acetylglucosamine-derived O-antigen building blocks biosynthesis (PWY-7332) and super pathway of thiamin diphosphate biosynthesis I (THISYN-PWY). We repeated this differential analysis using the ALDEx2 option provided in Namco. ALDEx2 applies CLR transformation to the raw counts to address potential biases introduced through compositionality. Following this approach, no significant KO terms were identified. We hypothesize that CLR transformation might increase the specificity in functional analysis at the cost of sensitivity, suggesting that users need to carefully reflect on their method choice when interpreting their results.

**Figure: 7.**
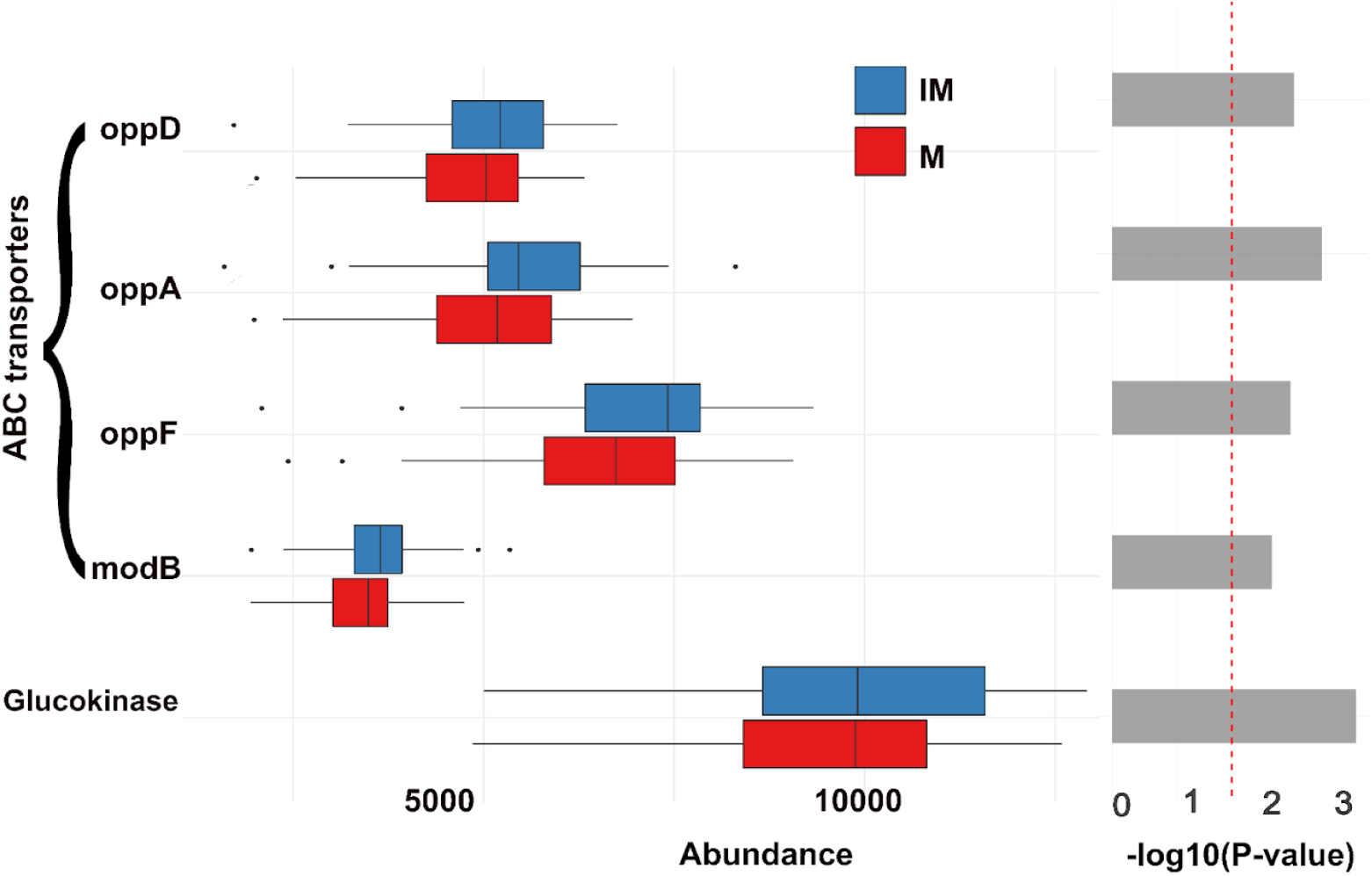
Bar plot showing the difference in the relative abundance of ABS transporters and glucokinase between IM and M group. p-values were calculated using the Wilcoxon Rank test on abundance values.

### Network Analysis

Co-occurrence networks were analyzed in Namco to characterize bacterial interactions between IM and M groups using the SPRING metric for network construction as the default option. To focus on the most abundant ASVs, an abundance cutoff of 0.25% and a prevalence cutoff of 10% were applied. The 270 remaining ASVs were used as input for network comparison. The co-occurrence network between IM and M at the genus level was illustrated in Figure 8. A genus-level network was generated using The SPRING (85) method as association measure (nlambda and replication numbers were set to 50 and 100, respectively). Eigenvector centrality was used for defining hubs and for scaling node sizes. Comparisons of all global measures for degree and eigenvector centrality, respectively, are given in Table 2. For none of the four centrality measures, any significant differences are observed. The two networks shared similar properties and no node hubs were identified in both groups. The largest differences in closeness centrality between IM and M groups belonged to the genus *CAG-56, Eubacterium coprostanoligenes*, *Lachnospira Lachnospiraceae UCG-004*, and *Ruminococcaceae Incertae Sedis*. Among these genera, only *Ruminococcaceae Incertae Sedis* was found to be significantly abundant in the IM group from previous analysis. Overall, there is no significant difference observed at the genus level network between the IM and M groups.

**Figure 8:**
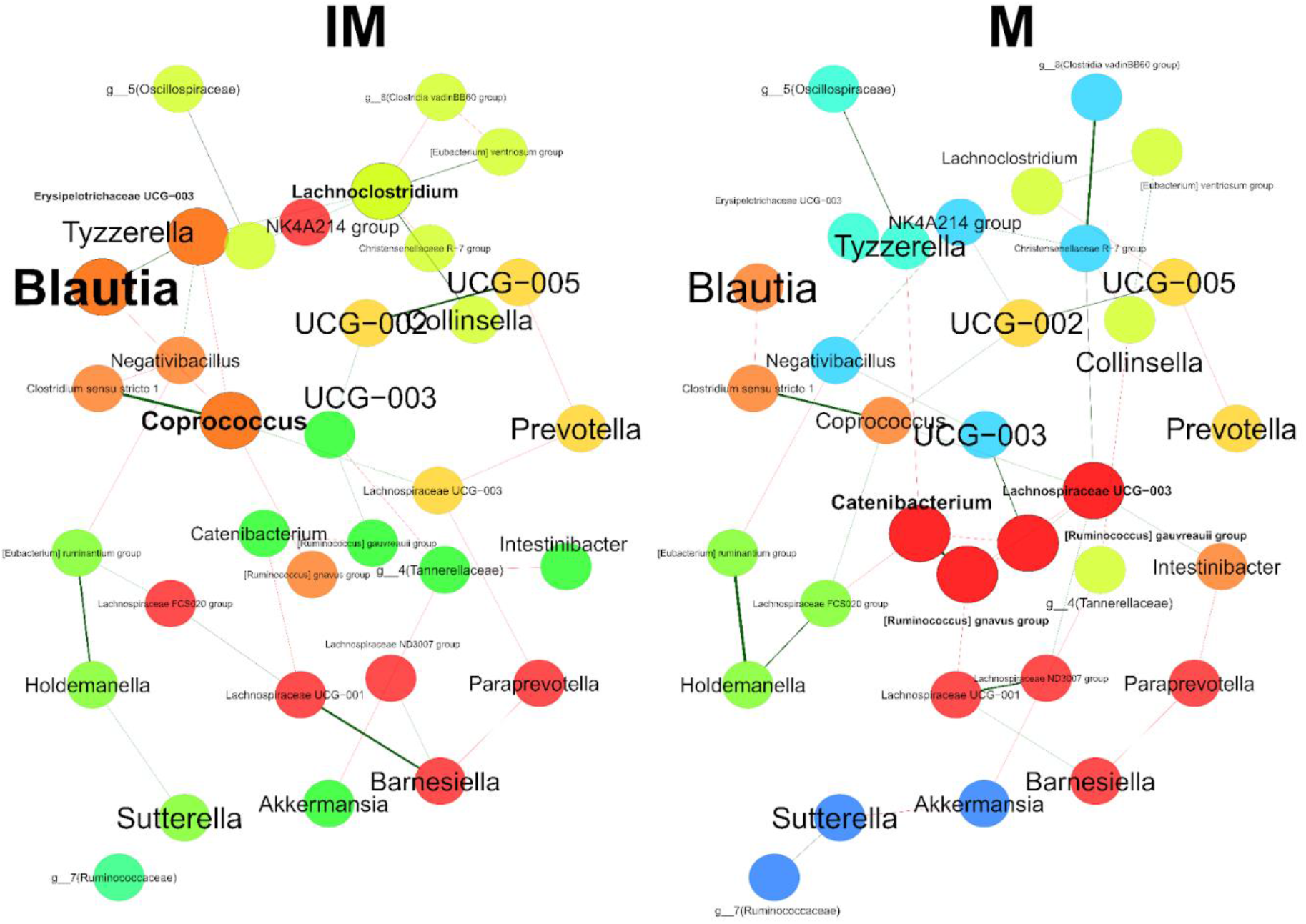
Bacterial associations on the genus level for the intervention (IM) and non intervention (M) groups using SPRING method. Green edges represent positive associations and red edges represent negative associations. Node colors represent clusters determined by using greedy modularity optimization. Networks are shown with only the 50 nodes with the highest degree and 50 edges with highest weight.

## Discussion

To gain biologically and clinically relevant insights from the large amounts of available microbiome sequencing data, a plethora of algorithms, statistical methods and software packages have been developed. We implemented Namco to serve as a one-stop data analysis platform that performs both raw data processing and basic as well as advanced downstream analyses of microbial datasets. Namco integrates previously available tools into a single coherent computational workflow and allows the user to construct, analyse and understand microbial composition in a fast and reproducible manner. This platform is intended to eliminate the use of command-line arguments during data processing. Namco is accessible in the web browser and hence does not require the installation of any software packages. Namco also allows saving results from each analysis as a R session which can be used to resume at any time, thus simplifying sharing of research results. Since Namco is available as a Docker image, it can be conveniently installed locally or on a clinical server behind a firewall to facilitate the GDPR-compliant analysis of sensitive data without the need for an upload to the public Namco instance. Detailed comparisons between other web-based tools (Table 1) showed that Namco offers a unique set of functions such as, for instance, time series clustering, function profiling using PICRUSt2, confounder analysis and topic modelling.

We considered a dietary intervention study as a case study to explore the features of Namco. We examined the association of rich fibre dietary intake with the gut microbiota composition through basic and advanced analysis in Namco. We investigated differences in relative abundance between IM and M groups, where we compared the top abundant taxa and also studied the intra-individual variation in gut microbiome with respect to fibre-enriched diets. After exploring the datasets in terms of relative abundance and diversity analysis, we found that genera with significant differences between IM and M groups were involved in the production of butyrate which is a SCFA that helps to maintain the homeostasis of the gut via anti-inflammatory and antimicrobial actions (86–88). Namco did not only provide information about differential abundant microbial composition but also helped in determining the significantly different KO terms and pathways. Namco revealed that the IM group showed a positive association with the presence of the glucokinase which belongs to amino sugar and nucleotide sugar metabolism. High fibre intake showed a positive impact on glucose metabolism in humans. Studies showed that long term intake of fibre improves the glucose homeostasis (89). In addition to that, Namco also identified ABC transporters which play a major role in the transmission of glucose through plasma membranes as significantly correlated with the IM group (82). It was also possible to study microbial interactions by generating differential microbial co-occurrence networks on the genus level using Namco. The topological features of the resulting differential network from IM and M groups showed only a slight difference in the estimated associations. Overall, Namco provided a much-needed interface to analyze the microbial community data in a more intuitive way.

## Conclusion

In summary, we present Namco, a shiny R application dedicated to providing end to end microbiome analysis for 16S rRNA gene analysis. We incorporate leading R packages for both upstream and downstream analysis in an efficient framework for researchers to characterize and understand the microbial community structure in their data, leading to valuable insights into the connection between the microbial community and phenotypes of interest. In the future, we plan to further expand Namco with support for novel analysis techniques and for correlation of microbial abundances with other data sources such as metabolomics and transcriptomics.

## Data summary

1. Namco was implemented as a R shiny app (https://exbio.wzw.tum.de/Namco/) under GNU General Public License v3.0.
2. The complete source code is publicly available at Github (https://github.com/biomedbigdata/Namco)
3. Sequence data used in the usercase was deposited in the National Center for Biotechnology Information (NCBI) SRA archive under the BioProject ID PRJNA774891.
4. User manuel is available at https://docs.google.com/document/d/1A_3oUV7xa7DRmPzZ-J-IIkk5m1b5bPxo59iF9BgBH7I/edit?usp=sharing

## Authors and contributors

**C**RediT taxonomy from CASRAI: **B.B., T.S., and H.H.; Investigation, Methodology, Resources, Validation. B.B., T.S., and H.H., A.D., M.M.; Data curation. A.D., M.Z., M.LA., and B.Ö.;** Software. **A.D., M.M.**; Formal analysis, **A.D., M.M;** Visualization, **M.M;** Writing – original draft. **M.LI., S.R.;** Conceptualization, Project administration, Supervision. **D.H., J.B, H.H.;** Funding acquisition. All authors were involved in Writing – review & editing.

## Consent for publication

Not applicable

## Acknowledgements

We also wanted to thank Luise Rauer and Claudia Hülpüsch from IEM Augsburg for constructive feedback during the development of Namco

## Repositories

1. Sequence data used in the usecase was deposited in the National Center for Biotechnology Information (NCBI) SRA archive under the BioProject ID PRJNA774891.

